# Infiltration of CD8^+^ T cells into tumor-cell clusters in Triple Negative Breast Cancer

**DOI:** 10.1101/430413

**Authors:** Xuefei Li, Tina Gruosso, Dongmei Zuo, Atilla Omeroglu, Sarkis Meterissian, Marie-Christine Guiot, Salazar Adam, Morag Park, Herbert Levine

**Author notes:** X.L. and T.G. contributed equally to this work. **Significance Statement** The infiltration of cytotoxic totoxic T cells into tumors is a critical factor in the efficacy of immunotherapy. Here we developed an algorithm to quantify the infiltration level using whole-section immunohistofluorescence (IHF) images from different patients. We observed consistent infiltration patterns of cytotoxic T cells (on the tumor-cell cluster level) at the core and margin of each tumor. These spatial distributions enabled us to generate several hypotheses for mechanisms underlying the various infiltration levels. We therefore used mathematical models to test which hypothetical mechanisms could recapitulate these patterns observed. One of the hypotheses, that of a fibrotic barrier, was shown to be unlikely. The other, involving a T cell chemorepellent, agrees with the data and can be tested in future experiments that directly measure the spatial pattern of chemokines.

## Abstract

Infiltration of CD8^+^ T lymphocytes into solid tumors is associated with good prognosis in various types of cancer, including Triple Negative Breast Cancers (TNBC). However, the mechanisms underlying different infiltration-levels are largely unknown. Here, we have characterized the spatial profile of CD8^+^ T cells around tumor-cell clusters in the core and margin regions in TNBC. Combining mathematical modeling and data analysis, we propose that there exists a possible chemo-repellent inside tumor-cell clusters, which prevents CD8^+^ T cells from infiltrating into tumor-cell clusters. Furthermore, investigation into the properties of collagen fibers suggests that variations in desmoplastic elements does not limit infiltration of CD8^+^ T lymphocytes into tumor-cell clusters. This is consistent with the prediction of our mathematical modeling analysis whereby CD8^+^ T cells are predicted to infiltrate the fibrotic barrier and their infiltration into tumor clusters is governed by other mechanisms involving a local repellent.

---

Activated CD8^+^ T lymphocytes have been demonstrated to be able to kill cancer cells via various mechanisms (1). Not-surprisingly, stronger infiltration of CD8^+^ T cells into tumors generally associates with better prognosis; this has been demonstrated in various cancer types such as melanoma (2, 3), ovarian (4), colorectal (5), bladder (6), breast (7), and pancreatic (8) cancer. Furthermore, stronger infiltration of CD8^+^ T cells can predict patient response to standard of care chemotherapy (9–11) and to immune checkpoint blockade therapy such as anti-CTLA-4 (12) or anti-PD-1 (13, 14). Therefore, it is important to characterize the infiltration of CD8^+^ T cells in solid tumors and mechanisms that regulate this.

Several efforts have been launched to quantify the distribution of CD8^+^ T cells at the whole-tumor level. For example, the “Immunoscore” was developed to evaluate the differences between the density of CD8^+^ T cells at the core (CT) versus the invasive margin (IM) of a tumor (15, 16). Promisingly, higher Immunoscore, essentially the ratio of T-cell density in CT over IM, is indicative of a good prognosis for colorectal cancer and melanoma patients (15, 17).

On the other hand, solid tumors usually consist of tumor-cell clusters interdigitated with non-tumoral (stromal) cells, which include T cells among other cell types. Within the tumor core, T cells can be constrained to lie within stromal regions in various types of cancer (18–22). The limited infiltration of CD8^+^ T cells into individual tumor-cell clusters is an indicator of worse prognosis (4, 23, 24) and lack of response to immune-blockade therapy (21, 25). Therefore, it is also important to quantify a complete spatial profile of CD8^+^ T cells at the tumor-cell clusters level and investigate possible mechanisms underlying differences in the spatial-infiltration patterns in different patients.

At least two mechanisms have previously been proposed to qualitatively explain the limited infiltration of CD8^+^ T cells into tumor-cell clusters: i) the physical-barrier hypothesis (26–29) and ii) the biochemical-barrier hypothesis (30, 31). In support of the physical-barrier hypothesis, CD8^+^ T cells were mostly observed to move back and forth along ECM (extra-cellular matrix) fibers that are parallel to the surface of tumor-cell clusters (29). Therefore, it might be difficult for CD8^+^ T cells to move across the fibers towards tumor-cell clusters. For the biochemical-barrier hypothesis, treating tumor spheroids (composed of both tumor cells and fibroblasts) with CXCL12 antibody can increase the number of infiltrating T lymphocytes (31).

In this paper, we focused on the infiltration profile of CD8^+^ T cells in Triple Negative Breast Cancers (TNBC) patient samples. TNBC represents 15-20% of all diagnosed breast cancers and lacks markers amenable to targeted therapies. Importantly, TNBC harbors heterogeneity in the level of immune infiltration and activation and furthermore the presence of tumor-infiltrated CD8^+^ T cells inside of tumor-cell clusters significantly reduces the relative risk of death from disease (24). Therefore, it is valuable to investigate mechanisms underlying different infiltration patterns in TNBC.

To evaluate whether a physical barrier or an alternative explanation such as a repellent barrier hypothesis could better explain the infiltration pattern of the CD8^+^ T cells, we developed a method to quantify the spatial profile of CD8^+^ T-cell density based on images containing immunohistochemical labelling of CD8^+^ T cells and cancer cells in TNBC patient whole-tumor samples. We quantified the infiltration pattern across both the tumor invasive margin as well as the boundary of tumor-cell clusters. By combining these spatial profile determinations with mathematical modeling studies and measurements of tissue fiber properties (such as length, density, thickness, alignment), our results strongly suggest that a physical barrier around tumor-cell clusters is not responsible for limiting the infiltration of CD8^+^ T cells into either the tumor as a whole or the tumor-cell clusters within. Instead, our results favor the hypothesis whereby biochemical factors such as the balance between chemoattractant and chemorepellent concentrations is the most likely mechanism underlying the observed CD8^+^ T-cell location profile. Our findings imply that different CD8^+^ T-cell profiles are a consequence of the different signaling properties of cancer cells in different patients.

## Results

### Spatial distribution of CD8^+^ T cells in primary tumors of TNBC

To gain an understanding of the factors impacting CD8^+^ T cell distribution patterns, the localization of CD8^+^ T cells was investigated and quantified following immunostaining with anti-CD8 (to label CD8^+^ T cells) and anti-pan-cytokeratin (to label epithelial tumor cells) in 28 whole-tumor TNBC specimens.

First, we assessed and quantified the infiltration patterns of CD8^+^ T cells across the tumor margin boundary (Fig. 1). For each specimen, we calculated the density of CD8^+^-pixels with respect to their distance from the tumor margin-boundary (Fig. 1). A 500*μm*-wide region centered on the margin boundary was defined as the “margin area” of a tumor (Region I, Fig. 1A). Another 500*μm*-wide region (250-750*μm* inside the margin boundary), was defined as the tumor core region proximal to the tumor margin boundary (Region II, Fig. 1A). For each specimen, we determined the infiltration level of CD8^+^ T cells as follows. The average density of CD8^+^ pixels in Region II (*< ρ_II_ >*) is compared to the maximum density in Region I 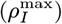 (see methods and Fig. 2). 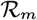 defined as 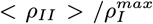. 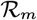 for all 28 patients and representative examples of profiles are presented in Fig. 2. Profiles of the CD8^+^-pixels are shown in Fig. S1. Based on their value of 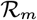, the tumors are divided into two groups: tumors where more CD8^+^ T cells accumulate in the margin area (called “margin-restricted”) versus tumors that have more CD8^+^ T cells infiltrated across the margin area (called “margin-infiltrated”). Representative examples for each group are shown in Fig. 2. Note that the cut-off values for separating different groups are being used just for illustrative purposes. The effect of ECM on the infiltration of CD8^+^ T cells, which will be shown in later sections, are evaluated by the correlation A ratio analysis and don’t depend on the assumed cutoffs.

We next performed a more detailed assessment of the tumor-cell clusters. For each sample, we selected 3 distinct regions within the tumor core (Fig. 1A-B) and quantified the density of CD8^+^-pixels with respect to their distance to the boundary of tumor-cell clusters (Fig. 1C). The tumor core is defined as at least 250 *μm* within the margin and the tumor-cell clusters are the tumor islands within the tumor core, separated by stroma (Fig. 1A). To estimate the infiltration level of CD8^+^ cells into tumor-cell clusters, we compare the average density of CD8^+^-pixel inside tumor-cell clusters (*< ρ_in_ >*) to the maximum density in the stroma 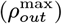 (see methods and Fig. 1C). Based on a ratio 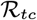 defined as 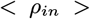 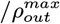, we further divided these tumors into 3 subgroups (see methods and Fig. 3A). The tumors in the first subgroup (6 patients) have 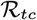 values (averaged by 3 different regions of the tumor core) below 0.05, and are termed “limited-infiltration” cases (for a representative example, see Fig. 3B). Tumors in the second subgroup (18 patients) with 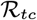 values between 0.05 and 0.5 are termed“intermediate-infiltration” cases (for a representative example see Fig. 3C, while tumors in the third subgroup (4 patients) with 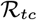 values above 0.5 are termed “full-infiltration” cases (for a representative example, see Fig. 3D). The spatial profiles of CD8^+^ T cells for all patients are depicted in Fig. S2. Note again that the cut-off values for separating different groups are being used just for illustrative purposes. In addition, it is important to note that in most tumors of the intermediate-infiltration subgroup, the CD8^+^ T cell density profile inside the tumor-cell clusters displays a specific feature: the density first decreases when moving from the boundary to the center of the tumor-cell clusters and then rises again when approaching the center (Fig. 3D).

### Mathematical models considering only reduced T-cell motility around tumor-cell clusters predicts T-cell profiles that are inconsistent with experimental observations

Next, we tested whether a mechanism-based model of T cell infiltration dynamics could explain the observed CD8^+^ T cell localization patterns. We have limited this attempt to the tumor-cluster level data, as there are clear consistent structural elements in these findings (for example, we observed a consistent accumulation of CD8^+^ T cells outside of tumor-cell clusters for all tumors with limited and intermediate CD8^+^ T cell infiltration, Fig. S2) as opposed to the ones related to the margin (for example, the peak of the CD8^+^-pixel density was not consistently located outside of the tumor margin, Fig. S1). We believe that this difference arises because the overall geometry of the tumor is highly patient-specific and this has a large effect on the ability to cross the margin.

It has been shown that the motility of CD8^+^ T cells can be reduced by ECM in human lung and ovarian tumors (19, 29), and this has been hypothesized to be responsible for the limited infiltration of CD8^+^ T cells into a solid tumor (27). Therefor we used a mathematical modeling approach to test the effects of reduced T-cell motility on the spatial distribution of T cells.

In all of our models, we assume that CD8^+^ T cells follow the gradient of a chemo-attractant and hence migrate toward a tumor-cell cluster. Stochasticity in the migration of a T cell is modeled by an effective Brownian diffusion. The possible reduced motility of CD8^+^ T cells in the matrix of collagen fibers is modeled by reduced effective diffusion and chemotaxis coefficients (see Methods).

**Fig. 1.**
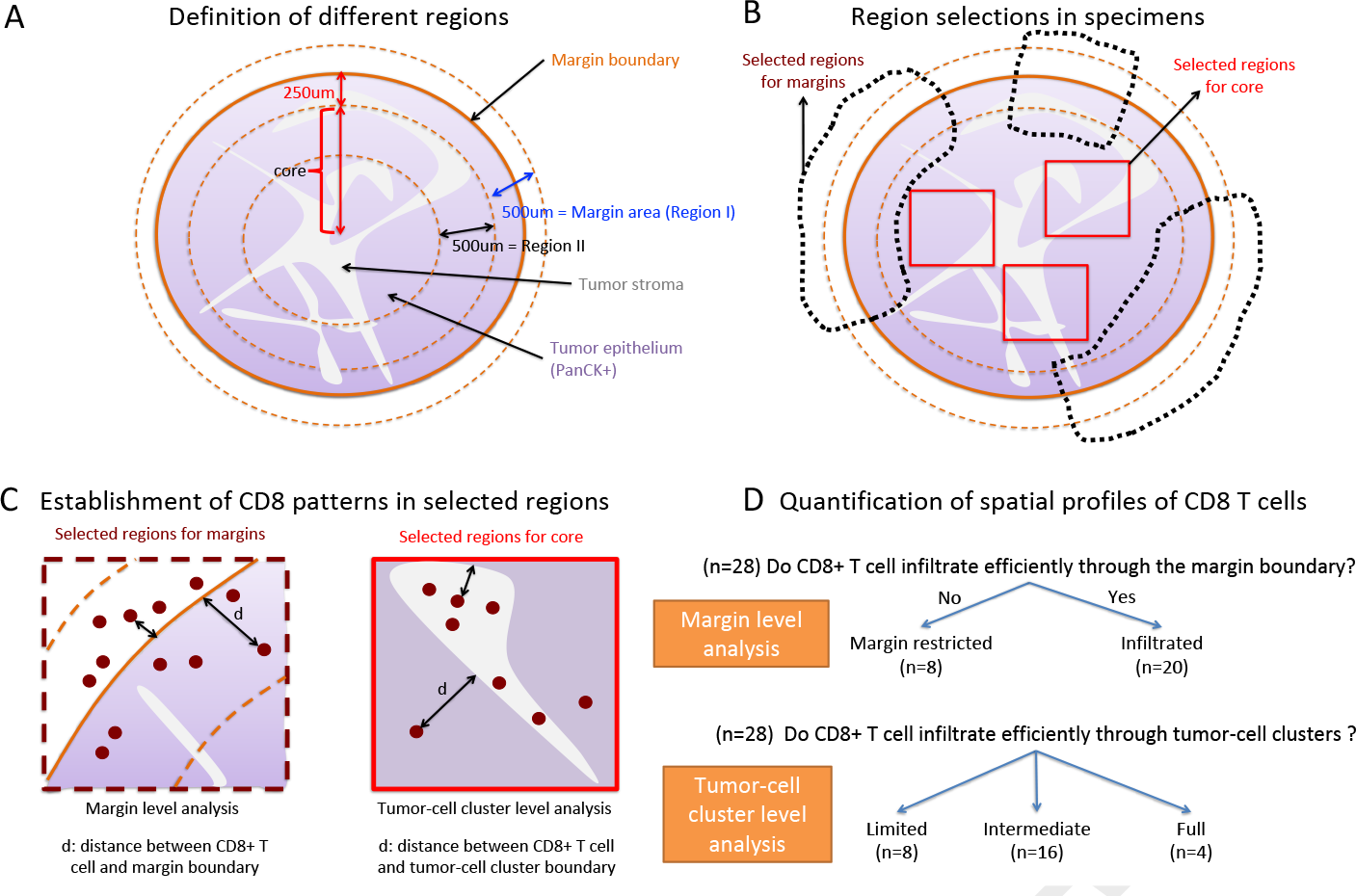
A. Illustration of different regions defined in the image analysis. B. Illustration of the selected regions of tumor margin and tumor core. For the tumor margin, parts of the margins were excluded whenever the juxtatumoral tissue did not exist. For the tumor core, whenever possible three regions (1.95*mm* × 1.95*mm*) were selected for each tumor. C. Illustration of the CD8^+^-T-cell-profile calculation for two levels: margin-boundary level and tumor-cell cluster level (see Methods). D. Based on the spatial profile of CD8^+^ T cells at the tumor margin and the tumor core, 28 patients were grouped into 2 groups (margin-boundary level) and 3 groups (tumor-cell cluster level), respectively.

**Fig. 2.**
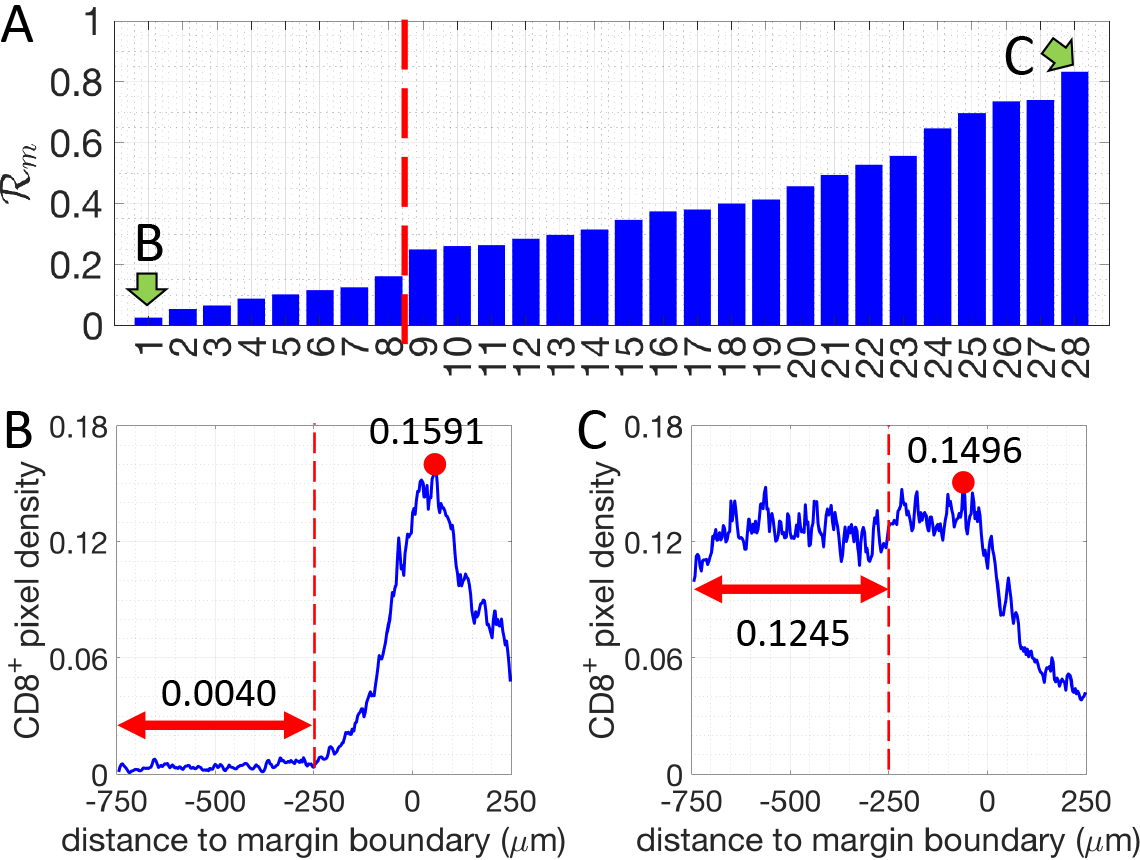
The density ratio of CD8^+^ pixels for all 28 patients analyzed (margin-boundary level analysis). Two patients, indicated by green arrows, are selected to illustrate the corresponding CD8^+^-pixel profiles. B and C. Examples of the density of CD8^+^ pixels (blue lines) for one margin-restricted tumor and one infiltrated tumor, respectively. The average densities in the region II (Fig. 1, between, −750*μm* and −250*μm*, red double-arrowed lines) are as indicated. The maximal densities (red dots) in the margin area (region I) (between −250*μm* and 250*μm*) are as indicated.

**Fig. 3.**
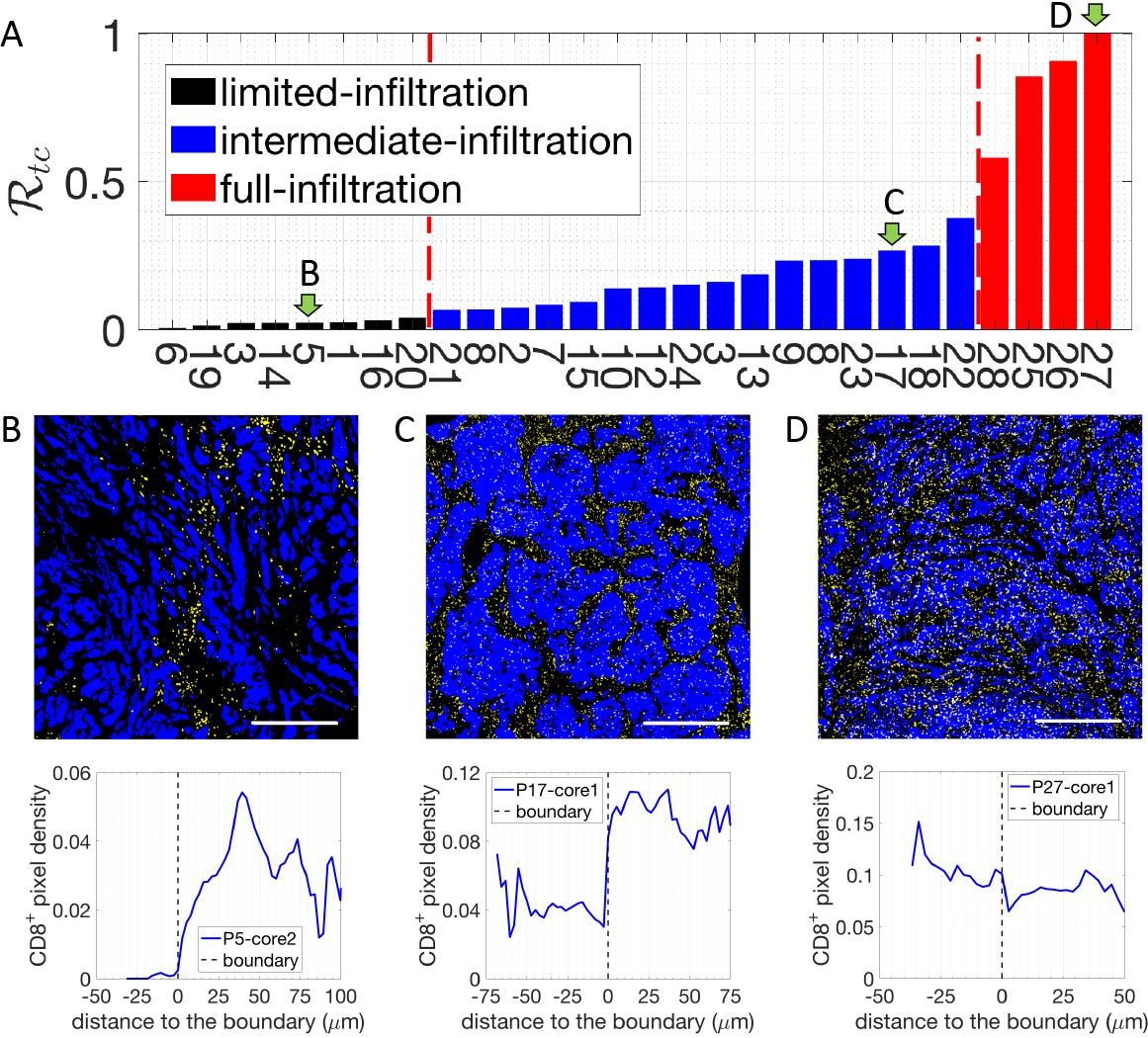
A. The density ratio of CD8^+^ pixels 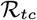 for all 28 patients. Three patients, indicated by green arrows, are selected to illustrate the corresponding CD8^+^-pixel profiles on the tumor-cell cluster level. B. Representative image with limited CD8^+^ T cell infiltration inside of tumor-cell clusters (up) and the corresponding CD8^+^ T cell distribution quantification (bottom). C. Representative image with intermediate CD8^+^ T cell infiltration inside of tumor-cell clusters (up) and the corresponding CD8^+^ T cell distribution quantification (bottom). D. Representative image with full CD8^+^ T cell infiltration inside of tumor-cell clusters (up) and the corresponding CD8^+^ T cell distribution quantification (bottom). In the representative images, tumor epithelial cells are in blue (Pan-Cytokeratin) and CD8^+^ T cells in yellow (CD8). Scale bar 500 *μ*m.

Initially, we tested an “abrogated T ceif motiiity” scenario, in which dense collagen fibers would directly limit T cell migration as soon as a T cell contacts such fibers outside the tumor-cell cluster. In this scenario, the density of CD8^+^ T cells would reach its maximum exactly at the boundary of the dense fiber region, and decay to lower values as we move through the stroma towards the tumor cluster (Fig. 4A). In the samples examined, CD8^+^ T cells were not found inside the dense fiber region. Thus, this predicted profile of CD8^+^ T cells does not match the experimental observations as shown in Fig. 3. Based on this analysis, an “abrogated T cell motility” scenario is not a likely mechanism for the observed T-cell infiltration patterns.

Next, we tested a “reduced T cell motility” scenario in which the motility of CD8^+^ T cells is reduced, but not completely abrogated, in the region with dense collagen fibers (see Methods). In this model, T-cell are still able to migrate toward the center of the tumor-cell cluster (Fig. 4B). Over time, the model predicts that the peak of T-cell density gradually moves into the tumor core and eventually most CD8^+^ T cells would occupy the center of the tumor-cell cluster instead of accumulating at the boundary. Therefore, this hypothesis would lead us to expect that in different patients the density of CD8^+^ T cells peaks would be located at various infiltration depths inside tumor-cell clusters. However, this scenario was not observed in the samples investigated, rendering it unlikely as well that this simplified “reduced T cell motility” scenario represents the main driver of CD8^+^ T cell infiltration patterns.

In the aforementioned “reduced T cell motility” scenario, CD8^+^ T cells maintain their reduced T cell motility as they infiltrate the tumor epithelium and thus display very limited movement inside tumor-cell cluster (Fig. 4B). We next modified this model by testing the role of a potential regaining of CD8^+^ T cell motility in the tumor-cell clusters as a consequence of a decreased fiber network density (“reduced followed by regained T cell motility” scenario) (Fig. 4C, spatial pattern of the diffusion coefficient is represented as a dash line). For example, T cell movement could be impeded by the dense collagen fibers in stroma, but would be regained as they progress within the tumor epithelium and move away from the dense fibers. In this scenario, CD8^+^ T cells will be attracted to tumor-cell clusters and transiently accumulate in the region with dense collagen fibers. Furthermore, if we assume that CD8^+^ T cells are no longer attracted to the center of the tumor-cell clusters once inside the clusters, in the intermediate time, CD8^+^ T cells will then diffuse within the tumor-cell cluster and their profile will monotonically decrease from the cluster boundary into the center of the cluster. In the long-time limit, CD8^+^ T cells would be distributed homogeneously inside tumor-cell clusters (Fig. 4C). At an early time in this “reduced followed by regained T cell motility” model, the modeled CD8^+^ T cell profile (red line in Fig. 4C) is qualitatively similar to the observed limited-infiltration profiles (Fig. 3A and Fig. S2). However, for the intermediate and full infiltration groups, only one third (12 out of 35) of the observed profiles (Fig. S2, intermediate-infiltration) are qualitatively similar to these predicted ones (blue, black and green lines in Fig. 4C). Indeed, in the other samples with an intermediate level of CD8^+^ T-cell infiltration, the observed profile is not monotonic inside tumor-cell clusters but rather displays an accumulation of CD8^+^ T cells in the center of tumor-cell clusters (Fig. 3B, Fig. S2). In conclusion, this “reduced followed by regained T cell motility” scenario can only explain a limited range of the observed the CD8^+^ T-cell infiltration profiles and only by assuming that these patterns are in a transient phase.

We further investigated other parameters and assumptions that could augment this “reduced followed by regained T cell motility” scenario. For example, we can allow the CD8^+^ T cells to also regain their chemotactic ability inside tumor-cell clusters (Fig. 4D). This scenario permits an accumulation of CD8^+^ T cells inside tumor-cell clusters but the steady-state shapes of the T-cell profiles again do not fully recapitulate the experimental ones shown in Fig. 3B. Specifically, in the long-time steady-state limit of this model, instead of reaching a homogeneous distribution, the modeled density of CD8^+^ T cells inside tumor-cell clusters decreases from the center to the boundary (Fig. 4D, solid green line). This distribution of CD8^+^ T cells is similar to most of fully infiltrated tumors but not of the others (Fig. 3C and Fig. S2). In addition, similar to the modeling results in Fig. 4C, at an early time, the modeled profile (red line in Fig. 4D) is also qualitatively similar to the observed limited-infiltration profiles (Fig. 3A and Fig. S2). In conclusion, a scenario with “reduced followed by regained T cell motility and chemotaxis” could explain the observed CD8^+^ T cell infiltration profile in limited and fully infiltrated TNBC tumors.

Therefore, using mathematical models including both a physical barrier term giving rise to a reduced followed by regained T cell motility and chemotaxis, we are able to explain the observed infiltration profile of CD8+ T cells in patients with limited or full infiltration of CD8^+^ T cells (Fig. S2). However, this hypothesis cannot robustly explain the CD8^+^ T cells profiles for many patients with an intermediate infiltration of CD8^+^ T cells. Furthermore, we need to appeal to transient profiles in this type of model, which means that eventually, we should expect the full infiltration of CD8^+^ T cells in the long-time limit for all patients. Based on the measurement of motility coefficient of T cells in a tumor nest by Salmon et al. (19), one T cell can cover the length of 100*μm* (the typical size of tumor-cell clusters observed in our samples) in 6 hours with a diffusion coefficient 5 *μm*^2^/min. Therefore, T cells should be able to fully infiltrate the tumor-cell clusters through diffusion within days, which is a very much shorter time scale than the clinical course of disease. However, patients with a full infiltration of CD8^+^ T cells are uncommon (only 4 out of 28). These observations thus lead us to consider alternative mechanisms.

### Mathematical models with a hypothetical repellent predicts T-cell profiles that resemble profiles observed in patient data

It is clear that biochemical signaling processes can play a role beyond purely biophysical ones in determining T-cell infiltration. For example, Lyford-Pike et al. (32) explicitly showed that PD-L1 is enriched at the boundary of tumor-cell clusters, which may create a biochemical immunoprotective “barrier” for tumor-cell clusters. More recently, the role of cytokine signaling by tumor cells has been shown to be critical for infiltration in a genetically engineered mouse pancreatic cancer system (33). We therefore tested a simple scenario involving a repulsive interaction between CD8^+^ T cells and cancer cells. Specifically, we hypothesized that i) there is a repellent, secreted by cancer cells that are close to the boundary of tumor-cell clusters, ii) this repellent drives exclusion of CD8^+^ T cells from the region with high repellent concentration, and iii) all cancer cells secrete a T-cell attractant. In our model, if CD8^+^ T cells strongly respond to the repellent, T cells would barely infiltrate into the tumor-cell clusters (black solid lines in Fig. 5). In the same model, if the effect of the repellent is moderate, CD8^+^ T cells can infiltrate into the tumor-cell clusters (blue solid lines in Fig. 5). In addition, depending on the profile of the repellent (determined by the source locations and the repellent diffusion coefficient) and the chemotactic ability of T cells, the modeled T-cell profile can be monotonic (Fig. 5A) or non-monotonic (Fig. 5B) inside tumor cell clusters. Lastly, if the effect of the repellent is very weak, T cells will infiltrate extensively into the tumor-cell clusters (red solid lines in Fig. 5). In this case, T cells distribute homogeneously inside the tumor-cell cluster (Fig. 5A) or accumulate in the center (Fig. 5B), depending on the profile of the repellent and the chemotactic ability of T cells in following the gradient of attractant. Notably, the profiles shown in Fig. 5 are all steady state distributions of CD8^+^ T cells. In conclusion, a scenario with both a repellent and an attractant of T cells could robustly explain all observed CD8^+^ T cell infiltration patterns in TNBC tumors.

### Desmoplastic elements are not limiting lymphocytic infiltration in TNBC

Our modeling results strongly suggest that a purely physical motility barrier cannot fully account for the variety of T cell infiltration pattern seen in our TNBC patients. We therefore studied in detail whether this prediction as applied to fibrosis (19, 34). Fibrosis is elevated in TNBC and HER2 tumors relative to other breast cancer subtypes (35)) as assessed though increased collagen cross-linking and thickening. We first assessed whether the stroma at the margin of tumors, where the infiltration of CD8^+^ T cells decreases, demonstrates increased collagen deposition and cross-linking as visualized by polarized-light imaging of Picrosirius Red-stained sections. We did detect thick collagen fibers at the tumor margin as compared to the tumor core, consistent with previously observations (35). However, no significant correlations were observed between the thickness of collagen fibers and the level of CD8^+^ cell margin infiltration. (Fig. 6A and 6B). To further characterize extracellular matrix (ECM) status, we performed Second Harmonic Generation (SHG) imaging using two-photon microscopy (Fig. 6C) that allowed us to quantify fiber density, fiber length and fiber alignment. The fiber alignment is defined by the Coefficient of Variation (CV) of the angle for all fibers per tumor. The smaller the CV is, the more aligned the fibers are. This analysis revealed that the fiber density, fiber length and fiber alignment at the margin of the TNBC tumor samples does not correlate with the level of CD8^+^ cell margin infiltration (Fig. 6D).

To test the presence of a purely physical motility barrier at the tumor-cell cluster level, we also performed the same type of measurements on collagen fibers in the tumor core (Fig. 6E-H). Again, no significant differences in the properties of collagen fibers were observed among tumors with various infiltration levels into the tumor-clusters (Fig. 6E-H).

In conclusion, our data on the ECM structure in TNBC thus support our theoretical modeling results that disfavor the hypothesis of a stromal physical barrier preventing CD8^+^ T cell infiltration. As has previously been suggested in PDAC (8), desmoplastic elements do not appear to be the critical factor limiting lymphocytic infiltration in TNBC neither at the core nor at the margin level.

**Fig. 4.**
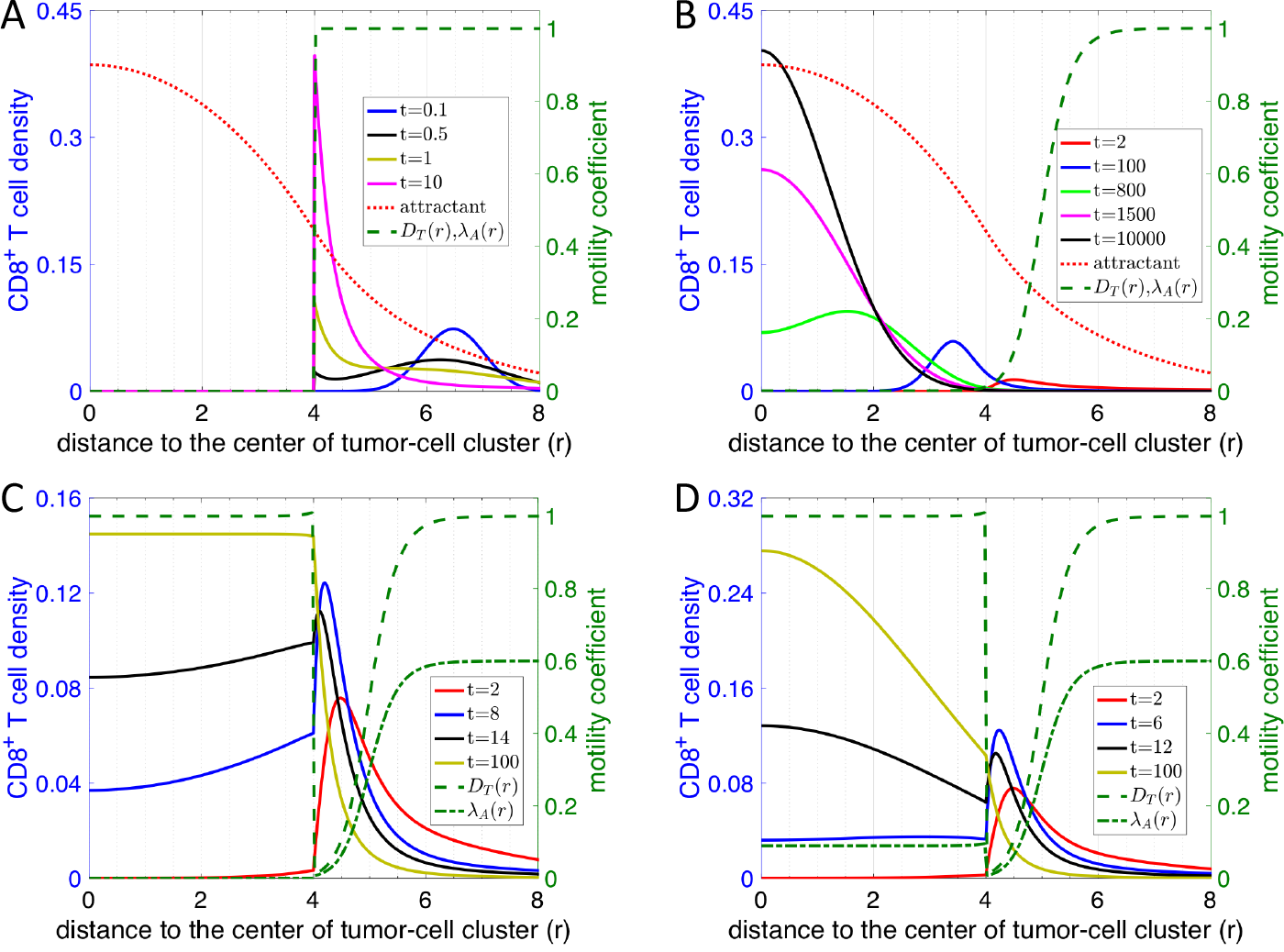
Time evolution of CD8^+^ T-cell profiles predicted by various scenarios of models based on the physical-barrier hypothesis. For all graphs, right y axis represents the diffusion (*D*_*T*_) and/or chemotaxis coefficient (λ_*A*_) of T cells as indicated in figure legend. Left y axis represents the attractant concentration and the CD8^+^ T cell distribution at different timepoints (in model unit) as indicated in the figure legend. The distribution of attractant is the same for all models. A. Abrogated T cell motility scenario where dense fiber prevent T cell infiltration. B. Reduced T cell motility scenario within regions of dense fibers and tumor-cell clusters. Noted that λ_*A*_ (*r*) is scaled by a factor of 1/30 in A and B. C. Reduced followed by regained T cell motility scenario where diffusion of T cells (*D*_*T*_) gradually decreases in the region with dense fibers but goes back to a normal level once T cells reach the region with tumor cells (dashed line). λ_*A*_(*r*) is scaled by a factor of 1/50 in the plot and is assumed to be zero once T cells move into tumor-cell clusters (dash-dotted line). D. Reduced followed by regained T cell motility and chemotaxis scenario. Scenario is similar to C with the exception that T cells retain their chemotaxis ability inside of tumor-cell clusters. λ_*A*_ (*r*) is scaled by a factor of 1/50 in the plot (dash-dotted line).

**Fig. 5.**
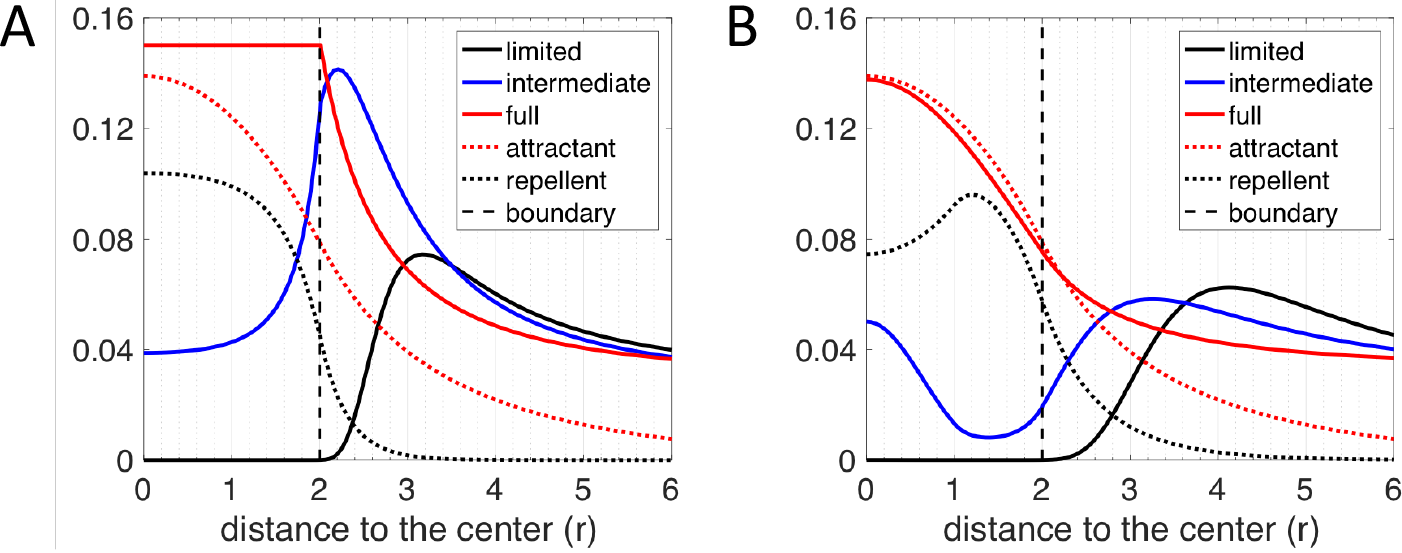
CD8^+^ T-cell profiles predicted by various scenarios of models based on the repellent-barrier hypothesis. For both graphs, y axis represents the long-term distribution of CD8^+^ T cells, attractant and repellent (scaled) concentration as indicated in figure legend. A. The chemotactic ability of CD8^+^ T cells in following the gradient of the attractant is assumed to be zero once T cells are inside of the tumor-cell cluster (*r* ≤ 2). The region of the source of the repellent is assumed to be between *r* = 0 and *r* = 2. Depending on how strongly the CD8^+^ T cells react to the repellent, the infiltration level can be limited (λ_*R*_ = 20, λ_*A*_ = 30), intermediate (λ_*R*_ = 20, λ_*R*_ = 30) or full (λ_*R*_ = 0, λ_*A*_ = 10). B. The chemotaxis ability of CD8^+^ T cells following the gradient of the attractant is assumed to be spatially uniform. The region of the source of the repellent is assumed to be between *r* = 1 and *r* = 2. Depending on how strongly the CD8^+^ T cells react to the gradient of the repellent, the infiltration level can be limited (λ_*R*_ = 40, λ_*A*_ = 60), intermediate (λ_*R*_ = 10, λ_*A*_ = 30) or full (λ_*R*_ = 0, λ_*A*_ = 10). The diffusion coefficient of repellent (*D*_*R*_ = 60) is assumed to 6 times larger than that in A (*D*_*R*_ = 10). Note that in A and B, the spatial profile of chemo-repellent is scaled by a factor of 0.1 and 0.2, respectively. the core nor at the margin level.

## Discussion

In this paper, we quantified the spatial distribution of CD8^+^ T cells with respect to their distance to both the tumor margin and the boundary of individual tumor-cell clusters. Generally, we observed that; i) patients differ both in the infiltration across the margin and infiltration at the cluster level; ii) for most samples, T cells mainly accumulate outside the tumor-cell clusters; iii) for some samples T cells can effectively infiltrate the tumor-cell clusters, and iv) for other samples, the T cell profile is intermediate. This last possibility reveals a nonmonotonic distribution of T cells, i.e., a drop of T-cell density at the boundary coupled with a second accumulation of T cells at the center of tumor-cell clusters.

Based on the quantified CD8^+^ T-cell profiles on the tumor cluster level, we constructed various mathematical models to test two hypothesized contributors affecting the spatial distribution of CD8^+^ T cells: a physical motility barrier set up by the ECM fibers in the stroma and a biochemical inhibitor ultimately due to the cancer cells inside the tumor-cell clusters. Mathematical models that only include physical barrier effects can qualitatively capture many (but not all) spatial features of the T-cell profiles. However, there is one significant shortcoming: the physical barrier scenario predicts that the profiles observed should be transient and hence eventually T cells should infiltrate all tumors. This appears inconsistent with simple time-scale estimates. Also, our data shows no correlation between cluster-level infiltration patterns and measures of typical ECM properties that could alter motility (Fig. 6). Furthermore, when observing the position of dense-fiber region and the region where lymphocytes accumulate at the tumor-margin, many lymphocytes can be found in the region close to the tumor-cell clusters whereas the dense-fiber region is further outside (Fig. S3). Our interpretation of this observation is that the dense fiber region might not be able to shield tumor cells from lymphocytes.

A biochemical model focusing on T-cell repulsion can give rise to the observed spatial profiles as steady-state solutions; the different patterns correspond to different properties of cancer cells in different patients. We therefore favor this modeling framework. Of course, a definitive test would require experiments which would enable us to study the infiltration of T cells in a time-dependent manner or alternatively perturb possible mechanisms with drugs to directly test their effects on the infiltration pattern of T cells.

We also investigated whether it is the stroma at the margin of the whole tumor that prevents the infiltration of CD8^+^ T cells on the whole tumor level. To do so we manually determined the outer boundary of tumors and quantified the spatial profile of CD8^+^ T cells with respect to this manually-curated boundary Although an accumulation of CD8^+^ T cells is still observed around the manually-curated tumor boundary for most patient samples, compared to the profiles quantified on the tumor-cell cluster level for the same region, the peak of CD8^+^ T cell density profile is not always outside of the outer boundary. From the actual images, this difference can be partially explained by the existence of T cells in the stroma between tumor-cell clusters but inside the tumor margin. This observation implies that: i) T cells can manage to get across the physical barrier if there is any; and ii) additional factors prevent T cells from further infiltrating the tumor. We have not tried to create an analogous mathematical model for these margin-related profiles, as one would clearly need to take into account the complex patient-specific geometry of the tumor and stroma. Nevertheless, our current repellent model offers a qualitative picture to understand why CD8^+^ T cells do not fully infiltrate the whole tumor: CD8^+^ T cells may have a difficult time to infiltrate tumor-cell clusters but still can move into the tumor core by following complex paths within the stroma.

For the biochemical “barrier” set up by the cancer cells, our current mathematical model utilizes an explicit diffusing repellent. In addition, in the model, the sources of the potential repellent are taken to lie at least partially inside of tumor cell clusters instead of being located exclusively at the boundary. This prediction could be verified once we have a good candidate for the repellent. Essentially, chemokines are the most likely candidates. For example, CXCL12, one of chemokines for CD8^+^ T lymphocytes, has recently shown to hinder the infiltration of T cells into spheroids formed by cancer cells and fibroblasts (31). Consistently, the gene-expression amplification of a chemokine cluster on chromosome 17 inversely correlates with the frequency of activated CD8^+^ T cells (36). A different possibly relies on the recent demonstration that TGF-*β* secreted by fibroblasts can contribute to exclusion of T cells at the margin (37); this might also happen on the tumor-cell cluster level. In addition, another recent analysis using gene expression data from TCGA database suggested that MAPK pathways are linked to reduced infiltration of cancer-fighting immune cells (38). Further investigation of the downstream secreted proteins of these MAPK pathways may reveal a possible candidate for the predicted repellent.

On the other hand, the interactions between other types of immune cells and CD8^+^ T cells can also be important for the final distribution of CD8^+^ T cells. For example, the exclusion of CD8^+^ T cells from tumors can be at least partially caused by the suppressed recruitment of one specific type of dendritic cell (CD103+) (39). Similarly the exclusion of T cells in a recent pancreatic cancer model could depend on myeloid suppressor cells (33). Therefore, investigating the spatial correlation of other types of immune cells with the CD8^+^ T cells may provide additional clues regarding the infiltration profile of CD8^+^ T cells.

In the current model, we did not consider the proliferation and death of T cells. There exists a possibility that those processes can contribute quantitatively to the spatial profiles of T cells. However, in order to explain the non-monotonic distribution of T cells inside the tumor-cell cluster, a complex spatially-dependent hypothesis on the cell birth/death rates would be required. We checked the expression of Ki67 in patient specimens using IHC. In the corresponding images, lymphocytes are identified as cells with smaller and round nucleus. Examples are summarized in Fig. S4. Qualitatively, most lymphocytes in the lymphocyte-cluster in stroma are not ki67+, which suggests that the accumulation of lymphocytes outside of tumor-cell clusters are mainly determined by te transport of lymphocytes. On the other hand, for patients belonging to the full-infiltration group, many lymphocytes inside tumor-cell clusters are also not ki67^+^, which suggests that those lymphocytes infiltrate into tumor-cell clusters via transport. Therefore, based on the current experimental observations and modeling analysis, we argue that the spatial dependent motility should be the major factor contributing to the observed CD8^+^ T cell profiles.

**Fig. 6.**
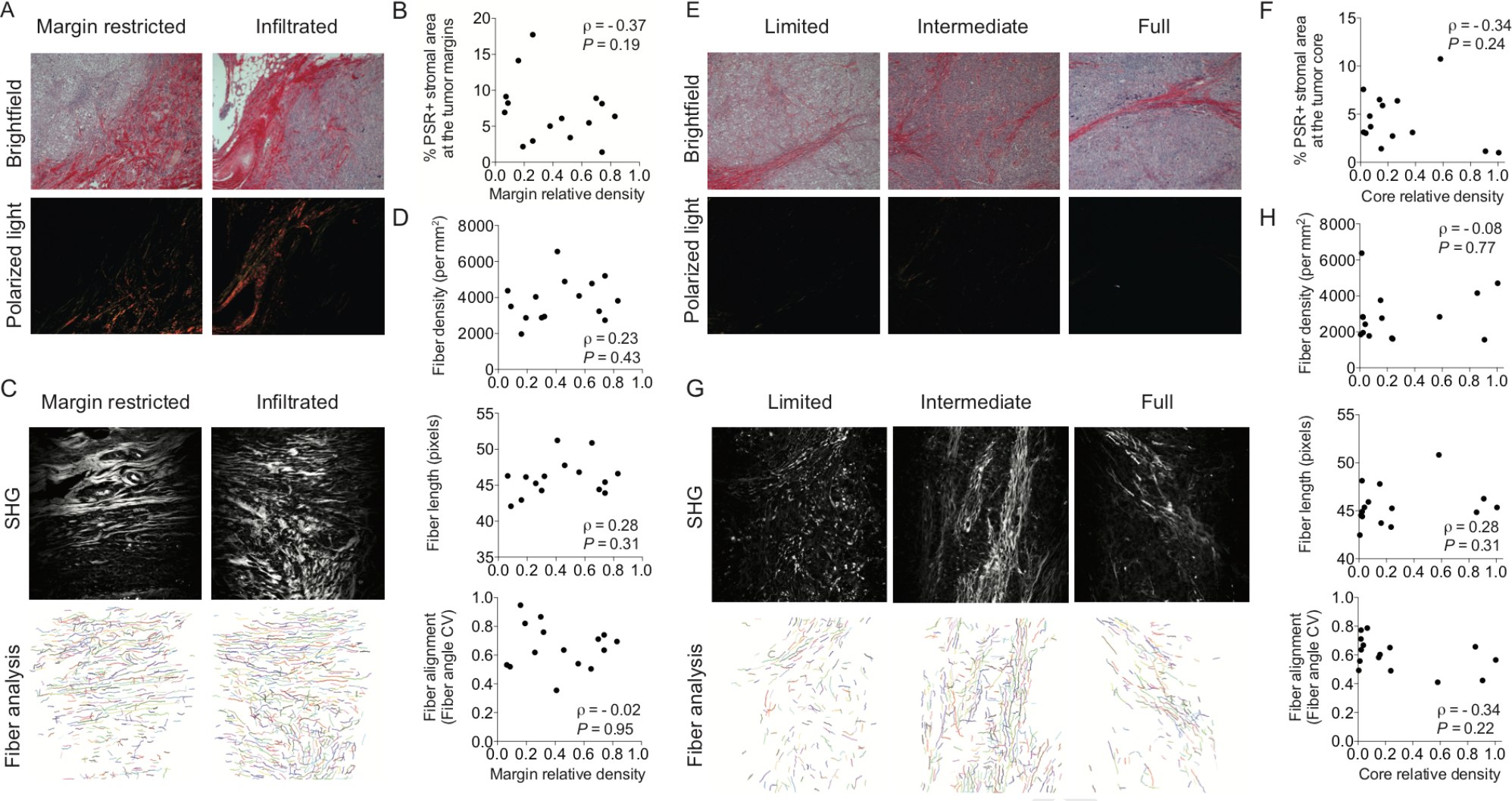
Desmoplastic elements are not limiting lymphocytic infiltration in TNBC. A. Representative images of Picrosirius Red staining at the tumor margins. Bright field images (upper panels) and matched polarized light images (lower panels) are presented. B. Quantification of Picrosirius Red polarized light signal in tumor margin areas. C. Second Harmonic generation images and representation of fiber individualization using the CT Fire software (tumor margins). D. Quantification of tumor margins fiber parameters as mentioned. E. Representative images of Picrosirius Red staining at the tumor core. Bright field images (upper panels) and matched polarized light images (lower panels) are presented. F. Quantification of Picrosirius Red polarized light signal in tumor core areas. G. Second Harmonic generation images and representation of fiber individualization using the CT Fire software (tumor margins). H. Quantification of of tumor margins fiber parameters as mentioned. Spearmann correlation

In summary, we studied the spatial-profile of CD8^+^ T cells around tumor-cell clusters and around the tumor as a whole, in TNBC patients. Combining data analysis and mathematical modeling, we provide evidence against the hypothesis that a physical barrier created by dense collagen fibers prevents the infiltration of CD8^+^ T cells into tumors and tumor-cell clusters. Instead, we propose that there could be a type of chemo-repellent inside tumor-cell clusters that prevents the infiltration. Further experiments on characterizing the level and patterns of chemokines inside tumor-cell clusters will be needed to verify the hypothesis in differing patient groups.

## Materials and Methods

### Calculation ofthe density profile of CD8^+^ T cells with respect to their distance to the tumor margin boundary

We developed an algorithm to extract the spatial information of CD8^+^ T cells from section images. For a given tissue section image, we manually drew the tumor margins. Next, a binary image of CD8^+^ pixels is generated by manually selecting a cut-off for the fluorescence intensity of CD8. With the spatial information of tumor-margins and each CD8^+^ pixels, the distance (d) between each CD8^+^ pixel and its nearest pixel on the tumor-margin boundary is calculated. The distance is binned and the size of the bin is 3-pixels (0.325*μm*x3). In addition, for all pixels, the nearest distance between each pixel and its nearest pixel on the boundaries is also calculated. Therefore, for each bin of the distance, the number of CD8^+^ pixels (*N*_*cD8*_(*d*) can be normalized by the total number of pixels (*N*_total_(*d*)), which gives the density of CD8^+^ pixels (*ρ*(*d*)) as a function of the distance to their nearest boundary pixels.

A ratio 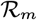 is defined to divide patients into 2 groups. The definition of 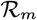 is given by the spatial average of *ρ* between *ρ* = −750*μm* and *d* = −250*μm* divided by the maximum of *ρ* between *d* = −250*μm* and *d* = 250*μm*. Here negative distance means the pixel is inside of tumor-margin boundary and vice versa.

### Calculation of the density profile of CD8^+^ T cells with respect to their distance to the boundary of tumor-cell clusters

For a given tissue section image, regions of tumor-cell clusters are identified by manually selecting a cut-off for the fluorescence intensity of pan-cytokeratin. Second, the boundary of tumor-cell clusters is identified using ImageJ based on the binary image of tumor-cell clusters. Next, a binary image of CD8^+^ pixels is generated by manually selecting a cut-off for the fluorescence intensity of CD8. With the spatial information of tumor-cell clusters and each CD8^+^ pixels, we then separate CD8^+^ pixels into two sets based on whether they overlap with tumor-cell clusters or not. For CD8^+^ pixels in each set, the distance (*d*_*in*_ and *d*_*out*_) between each CD8^+^ pixel and its nearest pixel on the boundaries is calculated. The distance is binned and the size of the bin is one pixel. At the same time, for all pixels, the nearest distance between each pixel and its nearest pixel on the boundaries is also calculated. Therefore, for each bin of the distance, the number of CD8^+^ pixels (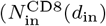 and 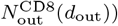 can be normalized by the total number of pixels 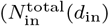 and 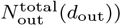, which gives the density of CD8^+^ pixels (*ρ*(*d*_*in*_) and *ρ*(*d*_*out*_)) as a function of the distance to their nearest boundary pixels.

A ratio 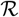 is defined to divide patients into 3 groups. The definition of 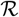 is given by the spatial average of *ρ*(*d*_*in*_) divided by the maximum of *ρ*(*d*_*out*_), i.e., 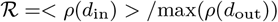.

### Mathematical modeling of the reduced T-cell motility by collagen fibers

In this type of model, we assume that CD8^+^ T cells follow the gradient of a chemokine (attractant) and migrate toward a tumor-cell cluster. The stochasticity of the migration of a T cell is modeled by an effective Brownian diffusion. The corresponding mathematical equations are as follows:

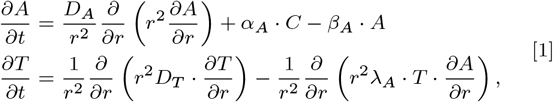

where *A* is the concentration of the chemotaxis attractant, *T* is the density of CD8^+^ T cells; *D*_*A*_ and *D*_*T*_ are the diffusion coefficient of the chemotaxis attractant and CD8^+^ T cells, respectively; /a is the chemotaxis coefficient of CD8^+^ T cells; α_*A*_ is the secretion rate of the chemotaxis attractant by tumor cells and β_*A*_ is the degradation rate of the attractant.

In our model, *D*_*A*_ is set to be 10. *D*_*T*_ and λ_*A*_ is a function of *r* (the distance from the center of a tumor-cell cluster). In Fig. 4A, *D*_*T*_ equals to 0 and 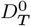 (= 1) inside and outside of the tumor-cell cluster, respectively. In Fig. 4B, 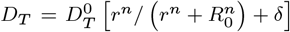 and 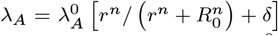. *R*_0_ = 5, *δ* = 0.001 and 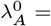 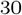. In Fig. 4C, *D*_*T*_ equals to 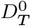 (=1) inside of the tumor-cell cluster. Outside of the tumor-cell cluster, 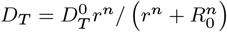. λ_A_ equals to 0 and 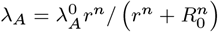 inside and outside of the tumor-cell cluster, respectively. In Fig. 4D, *D*_*T*_ is the same as that in Fig. 4C, whereas λ_A_ equals to 0.15×λ^0^_*A*_ 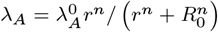 inside and outside of the tumor-cell cluster, respectively. *α*_*A*_ and *β*_*A*_ = 0.8 in Fig. 4. The density of tumor cells *C* equals to 1 and 0 for *r* < = 4 and *r* > 4, respectively. The boundary condition to solve Eq. [1] is (∂ *A*/∂*r*)|_r=0_, (∂*T*/∂*r*)|_*r* = 0_ = 0, *A*|_*r* = 10_ = 0, and (∂*T* / ∂ *r*)|_*r* = 10_ = 0.

### Mathematical modellng of the effects of a hypothesized repellent

In this type of model, the basic assumptions are the same as above: CD8^+^ T cells follow the gradient of a chemokine (attractant) and migrate toward a tumor-cell cluster and the stochasticity of the migration of a T cell is modeled by an effective Brownian diffusion. Furthermore, there exists a region inside a tumor-cell cluster where the CD8^+^ T-cell repellent is secreted. The corresponding mathematical equations are as follows:

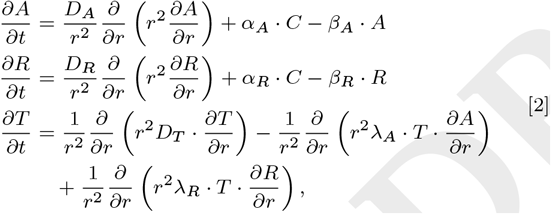

where *R* is the concentration of the chemotaxis repellent; *D*_*R*_ is the diffusion coefficient of the chemotaxis repellent; *λ*_*R*_ is the chemotaxis coefficient of CD8^+^ T cells with respect to the repellent *R*; *α*_*R*_ and *β*_*R*_ are the secretion and degradation rate of the chemotaxis repellent, respectively.

Here, *D*_*R*_ (= 10) and *D*_*T*_ (= 1) are constants. For different T-cell profiles in Fig. 5, the values of *λ*_*R*_ and *λ*_*A*_ are described in the figure caption fro Fig. 4. *α*_*R*_ = 85, *β*_*R*_ = 80. The density of tumor cells *C* equals to 1 and 0 for *r* <= 2 and *r* > 2, respectively. The boundary condition to solve Eq. [2] is (∂*A*/∂*r*)|_*r*=0_, (∂*R*/∂*r*)|_*r*=0_, (∂*T*/∂*r*)|_*r*=0_ = 0, *A*|_*r*=10_ = 0, *R*|_*r*=10_ = 0 and (∂*T*/∂*r*)|_*r*=10_ = 0.

### Sample collection and selection

Samples were collected from patients undergoing breast surgeries at the McGill University Health Centre (MUHC) between 1999 and 2012 who provided written, informed consent (MUHC REB protocols SDR-99-780 and SDR-00-966). All tissues were snap-frozen in O.C.T. Tissue-Teck Counpound within 30 minutes of removal. For the purposes of this study, samples were selected according to the following criteria: therapy-naive at time of surgical excision, clinically documented lack of expression/amplification of ER, PR and HER2, a histological subtype assignment of invasive ductal carcinoma (not otherwise specified) (IDC (NOS)) and availability of matched formalin-fixed paraffin-embedded (FFPE) tumor blocks. Information regarding clinical variables and disease course (follow-up) was obtained through review of Medical Records at the MUHC. 5*μ*m sections from frozen tissue were prepared for each sample, subjected to routine hematoxylin and eosin (H&E) staining, and evaluated by an attending clinical pathologist with expertise in breast tissue to identify invasive, in situ and normal components.

### Primary antibodies for Immunohistochemistry (IHC) and immuno-histofluorescence (IHF). See Table 1

#### IHC protocol

Sections were deparaffinized, conditioned and antigens were retrieved using proprietary buffers (pH6 or pH9). Slides were blocked for 5 minutes with Power Block reagent. Primary antibodies were applied at optimized concentrations overnight at 4°C, followed by 30 minutes of incubation with SignalStain Boost (Cell Signaling). Detection was performed with a DAB substrate kit (Cell Signaling). Slides were counterstained with Harris Hematoxylin.

**Table 1.** Primary antibodies

| Ab | Target Clone | Reference | Company | Dilution | Application | Antigen retrieval |
| --- | --- | --- | --- | --- | --- | --- |
| Ki67 | 30-9 | 790-4286 | Roche | 1 (prediluted) | IHC | EDTA |
| CD8a | C8/144B | M710301-2 | DAKO | 1/50 | IHF | Citrate |
| Pan-cytokeratin | AE1/AE3 & PCK26 | 760-2135 | Ventana | 1/2 | IHF | Citrate |

#### IHFprotocol

Sections were deparaffinized, conditioned and antigens were retrieved using proprietary buffers (pH6 or pH9). Slides were blocked for 5 minutes with Power Block reagent. Primary antibodies were applied at optimized concentrations overnight at 4°C, followed by 30 minutes of incubation with SignalStain Boost (Cell Signaling). Detection was performed with Tyramide Signal Amplification (TSA) kits (Thermo Fisher Scientific). Slides were counterstained with DAPI.

#### Picrosirius Red (PSR) Microscopyand imaging

FFPE tumor tissue tissues were sectioned and stained using 0.1% Picrosirius Red (Direct Red 80, Sigma) and counterstained with Weigert’s Hematoxylin, as previously described2. PSR polarized imaging was performed on an inverted Axio Observer Z1 microscope (Carl Zeiss Canada) using a 10X objective (PLAN NEOFLUAR, NA0.30, Ph 1) with linear polarizers and a quarter-wave plate. Halogen lamp intensity was kept constant for both image types, and an exposure time that optimized the signal-to-noise ratio was chosen and kept constant within each image type. All images were digitally captured using Luminara’s INFINITY3 color CCD camera (Cat No INFINITY3-6URC). 10 images were taken for each tumor (5 at tumor margins and 5 within tumor core). Images were quantified using ImageJ. Briefly, a minimal intensity threshold was used to eliminate the background and then the fiber density was measured as %stromal area covered by fibers in each image. This was performed on the whole image for margin images; for intratumoral stroma, stromal areas were manually identified from matched brightfield images and delineated prior to quantification.

#### Second Harmonic Generation (SHG) Microscopy and imaging

H&E sections were imaged using a Zeiss Axioexaminer upright microscope (Carl Zeiss Cananda). SHG signal was generated using 830nm excitation light from a Chameleon (Coherent) laser using a 20X objective (PLAN APOCHROMAT, NA0.8, DIC). All images were acquired with PMT detectors and using ZEN black software with consistent parameters. 6 images were taken for each tumor (3 at the tumor margins and 3 within the tumor core). Fiber individualization and quantitation of fiber parameters was performed using CT Fire software (freely available at http://loci.wisc.edu/software/ctfire)

## ACKNOWLEDGMENTS

This work was supported by the National Science Foundation Center for Theoretical Biological Physics (NSF PHY-1427654), NSF DMS-1361411, and The V Foundation. We are grateful to C. Moraes for his help in the analysis of fiber analysis as well as I. Acerbi, I. Dean and V. Weaver for their help with Picrosirius Red staining and analysis. We also thank Indica Labs and especially A. Hellebust for help with IHF analysis using HALOTM software. We are grateful to B. Clavieri (Microscopy Imaging Lab, University of Toronto) for scanning IHF slides. We thank Jo-Ann Bader and the staff at the Histology Core Facility at the Goodman Cancer Research Centre for assistance with sample preparation. We thank R. Deagle and E. Tse-Luen (Advanced BioImaging Facility, McGill University) for help with sample imaging. We thank members of the Depts of Surgery, Pathology and Anaesthesia at the McGill University Health Centre for their assistance with sample collection. We thank N. Bertos for help with the paper writing. This study was supported by funding from CQDM(Consortium québécois sur la decouverte du medicament/Quebec Consortium for Drug Discovery) and the NIH (National Institutes of Health) (to M.P.). This study was also supported by Merck, Sharpe & Dohme Corp./McGill Faculty of Medicine Grants for Translational Research (to M.P.). The breast tissue and data bank at McGill University is supported by funding from the Database and Tissue Bank Axis of the Réseau de Recherche en Cancer of the Fonds de Recherche du Québec-Santé and the Québec Breast Cancer Foundation (to M.P.). T.G. has been supported by the Charlotte and Leo Karassik Oncology fellowship.

---

**Fig. S1:**
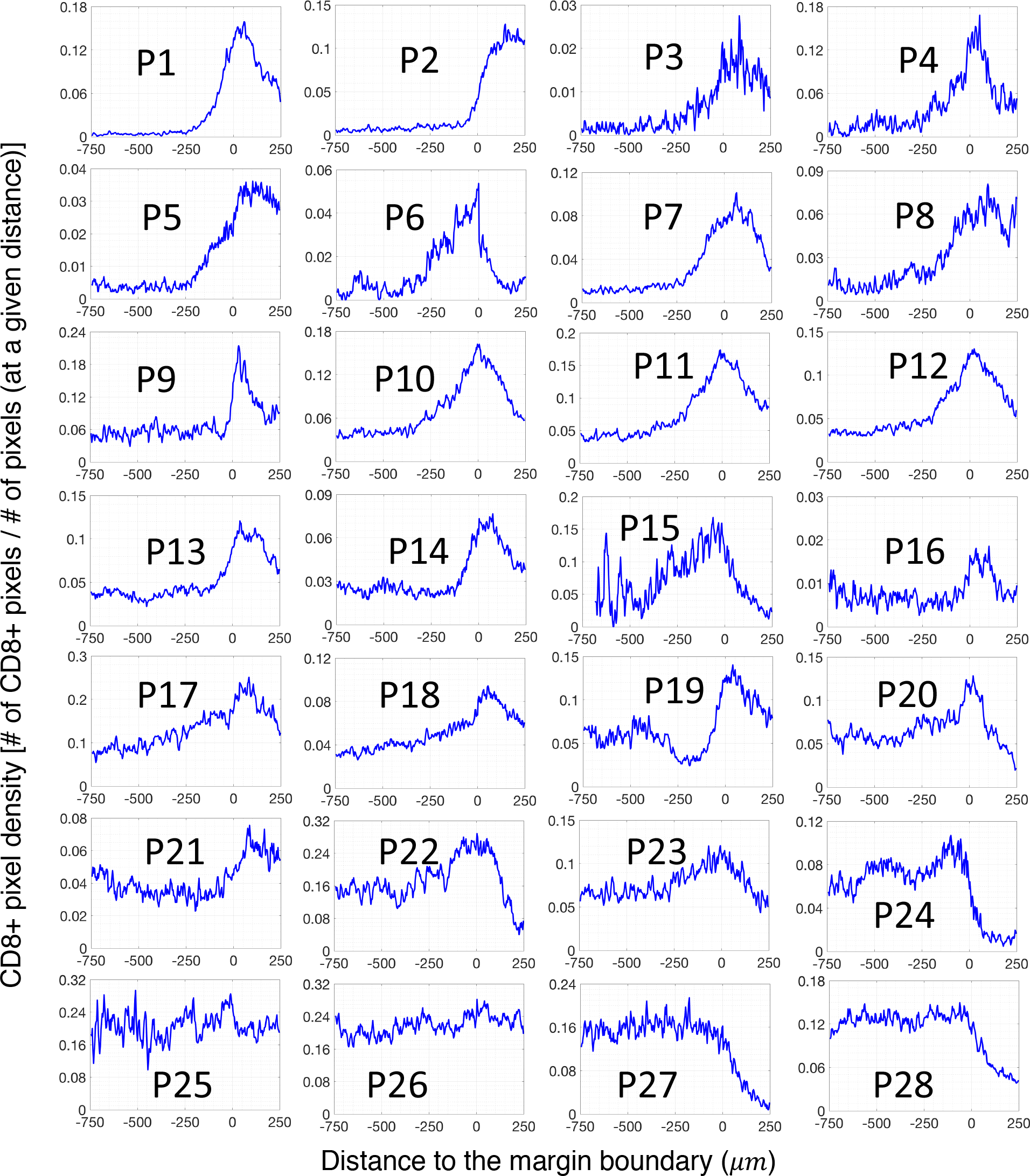
The spatial profile of CD8^+^-pixel density across the tumor-margin boundary for all 28 patients. We focused on the region between −750*μm* and 250*μm*around the margin boundary. The negative distance corresponds to the region inside of the margin boundary and vice versa.

**Fig. S2:**
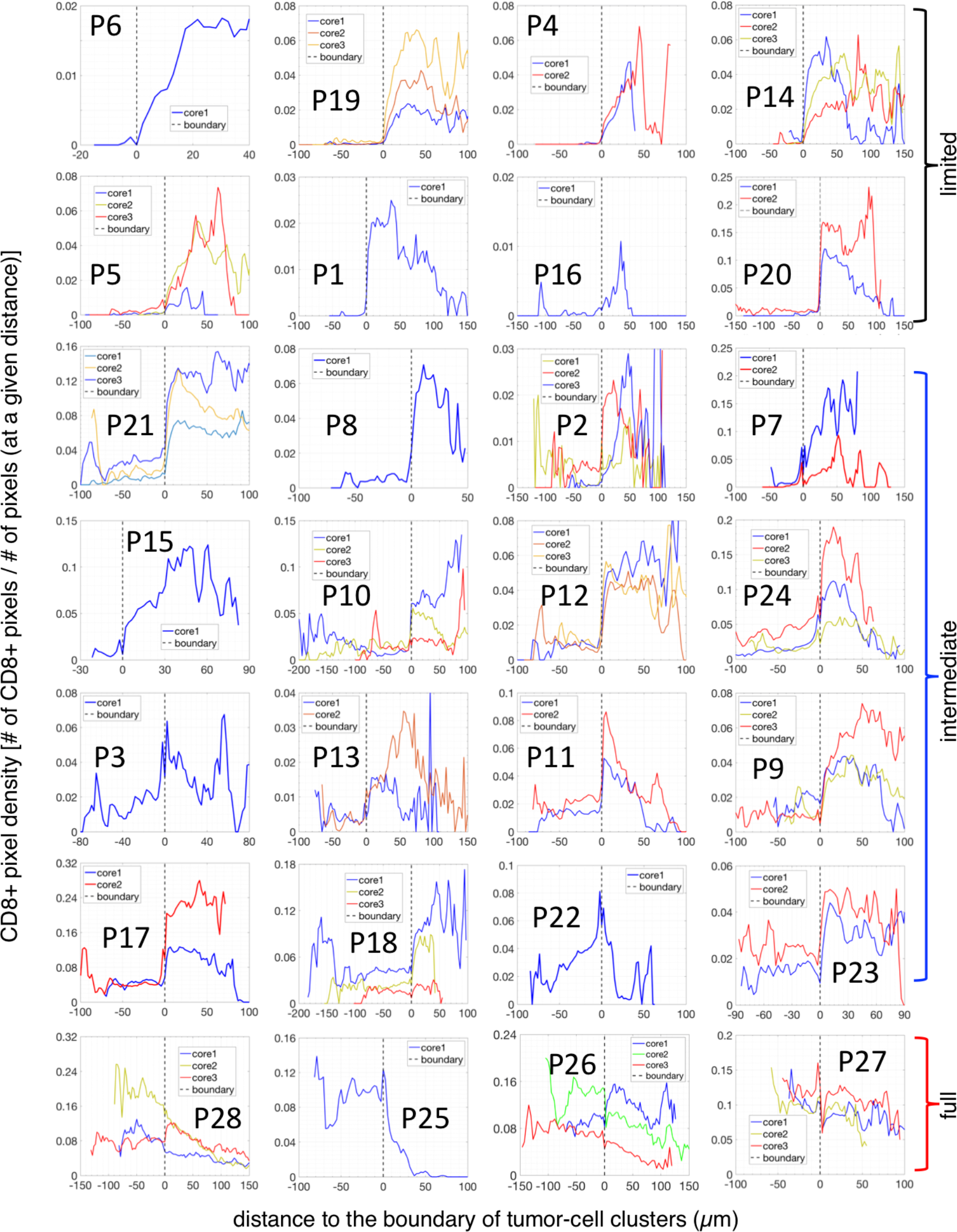
The spatial profile of the CD8^+^-density across the boundary of tumor-cell clusters for all 28 patients. The negative distance corresponds to the region inside of tumor-cell clusters and the positive distance corresponds to the stroma region in the tumor core.

**Fig. S3:**
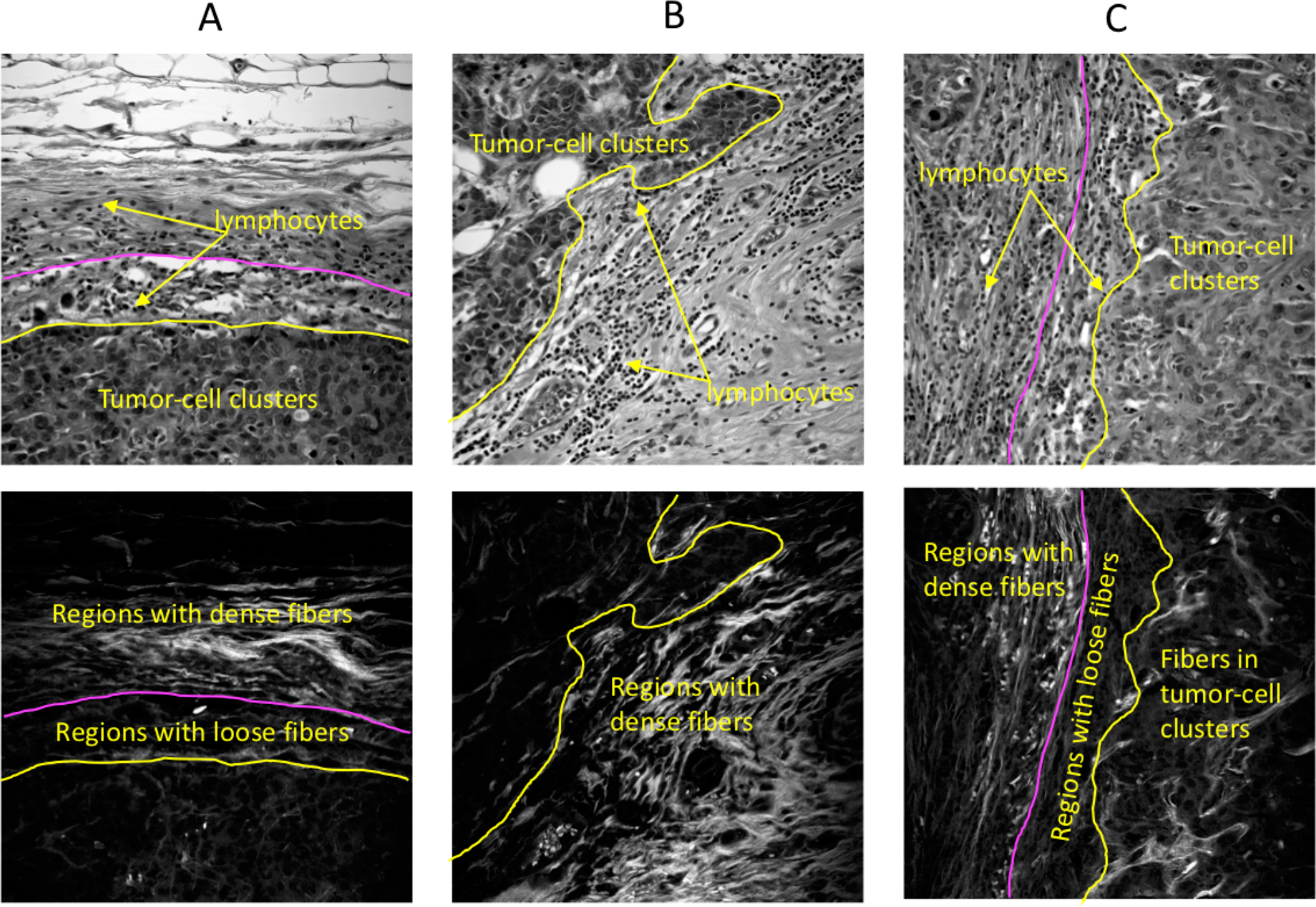
Representative invasive margins of three tumors are presented. Top row: phase contrast image of selected margins. Bottom row: Second-Harmonic Generation (SHG) images of the corresponding regions, which shows the pattern of collagen fibers (bright area). These examples show that lymphocytes can accumulate in regions close to the boundary of tumor-cell clusters instead of being constrained in regions with dense fibers. Representative lymphocytes are indicated by yellow arrows. Margins of the tumors are marked by yellow lines and the boundary between dense- and loose-fiber regions is manually marked by cyan lines.

**Fig. S4:**
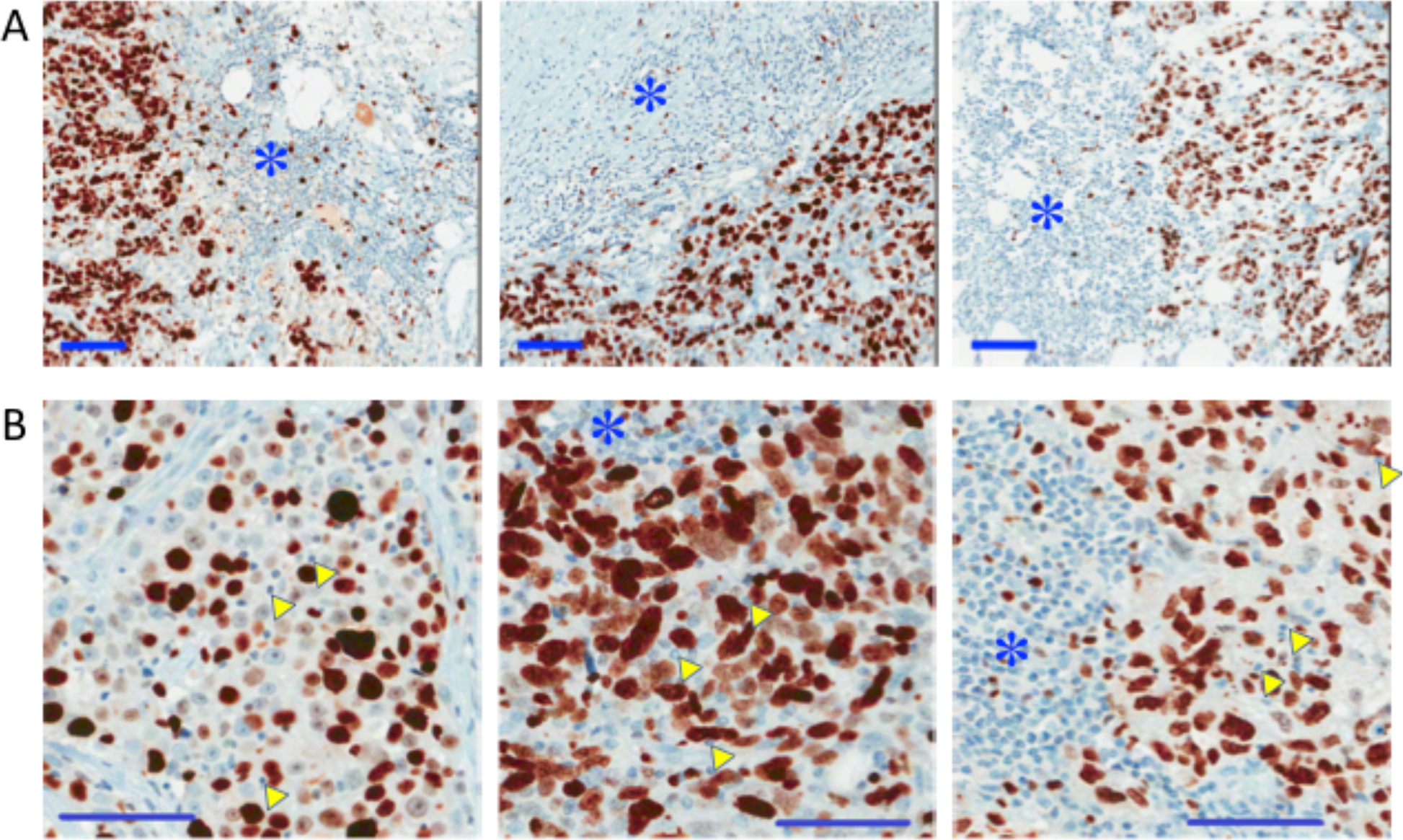
Examples of patient specimens with ki67 staining. A. These three examples are from the patient group with limited infiltration of CD8^+^ T lymphocytes, i.e., CD8^+^ cells are mostly outside of tumor-cell clusters. The three examples are from three different patients. In stroma (areas indicated with blue stars), most lymphocytes, characterized by small and round nucleus, are not ki67^+^. B. These three examples are from the patient group with full infiltration of CD8^+^ T lymphocytes, i.e., CD8^+^ cells density inside of tumor-cell clusters is higher than that outside. The three examples are from three different patients. Again, similar to the case in A, in stroma (areas indicated with blue stars), most lymphocytes are not ki67^+^. Furthermore, inside of tumor-cell clusters, many lymphocytes are not ki67^+^. Three lymphocytes are selected for each images for illustration, marked by yellow arrowheads. Scale bars: 100*μm*.

